# Protein structure prediction in the era of AI: challenges and limitations when applying to *in-silico* force spectroscopy

**DOI:** 10.1101/2022.06.30.498329

**Authors:** Priscila S. F. C. Gomes, Diego E. B. Gomes, Rafael C. Bernardi

**Affiliations:** Department of Physics, College of Sciences and Mathematics, Auburn University, Auburn, AL, 36849

**Keywords:** artificial intelligence, protein folding, adhesins, staph infection, molecular dynamics, force spectroscopy

## Abstract

Mechanoactive proteins are essential for a myriad of physiological and pathological processes. Guided by the advances in single-molecule force spectroscopy (SMFS), we have reached a molecular-level understanding of how several mechanoactive proteins respond to mechanical forces. However, even SMFS has its limitations, including the lack of detailed structural information during force-loading experiments. That is where molecular dynamics (MD) methods shine, bringing atomistic details with femtosecond time-resolution. However, MD heavily relies on the availability of high-resolution structures, which is not available for most proteins. For instance, the Protein Data Bank currently has 192K structures deposited, against 231M protein sequences available on Uniprot. But many are betting that this gap might become much smaller soon. Over the past year, the AI-based AlphaFold created a buzz on the structural biology field by being able to, for the first time, predict near-native protein folds from their sequences. For some, AlphaFold is causing the merge of structural biology with bioinformatics. In this perspective, using an *in silico* SMFS approach, we investigate how reliable AlphaFold structure predictions are to investigate mechanical properties of staph bacteria adhesins proteins. Our results show that AlphaFold produce extremally reliable protein folds, but in many cases is unable to predict high-resolution protein complexes accurately. Nonetheless, the results show that AlphaFold can revolutionize the investigation of these proteins, particularly by allowing high-throughput scanning of protein structures. Meanwhile, we show that the AlphaFold results need to be validated and should not be employed blindly, with the risk of obtaining an erroneous protein mechanism.

## 1 Introduction

Over the past year, the artificial intelligence (AI)-based software AlphaFold created a buzz on the structural biology field. For the first time, a software was able to predict near-native protein folds from their genetic sequence ^1^. DeepMind’s AlphaFold transformed, in principle, the protein structure solving problem that has been around for the past 50 years into a trivial task. The number of research papers and preprints citing the method soared since its code was released in July 2021 ^2^, with the accompanying article achieving about 1,000 citations (according to Google Scholar) in its first year.

The success of AlphaFold, and the analog RoseTTAFold approach ^3^ that appeared a few months later, is partially due to their open-source nature, which makes them readily and freely available to anyone one who is interested in trying these software. Furthermore, by pairing it with the European Bioinformatics Institute (EBI), AlphaFold has taken structural biology to the next level, allowing big consortiums to perform protein structure prediction to entire genomes, including human, mouse, *Saccharomyces* and *E. coli* ^4^. The resulting structures were made available on a database maintained by the EBI, containing almost a million structures: https://alphafold.ebi.ac.uk.

The broad spread use of AI-based structure prediction leads us to ask the question: How reliable are the structures predicted by such models? Despite the growing number of success stories ^1,5–9^, researchers are accumulating evidence showing that AI-based structure prediction methods are still nor perfect ^10,11^, and that there is ample room for improvement. In other words, some results suggest that both AlphaFold and RoseTTAFold are qualitatively great, but in many cases, they lack the level of details that is important to understand a protein function ^2,12,13^.

High-resolution protein structures are also crucial for drug-discovery. The ability to readily access the structure of any protein of the human genome is very attractive to those developing new drug compounds. Using an AI-based tool to predict how drugs bind to these proteins is an even larger challenge that will probably not be overcome soon due to the limited publicly available data for small molecule binding ^14^. In addition to that, AlphaFold lacks the precision to predict structural changes in consequence of mutations ^15^.

Working as a “computational microscope” molecular dynamics (MD) simulations are a unique tool to investigate biomolecules’ behavior with atomic resolution ^16–18^. However, as most computational chemistry methods, the quality of MD results relies heavily on the quality of the initial biomolecule structure ^19–21^. If AI-based structure prediction software are able to predict protein folds to the atomic level, MD simulations should be able to profit from these structures and give similar results to those obtained with experimentally determined structures.

A particularly powerful way of using MD simulations is by using it hand-in-hand with experimental methods. In the past few years, taking advantage of steered MD protocols, our group has pioneered what we call *in silico* single-molecule force spectroscopy (*in silico* SMFS) ^22–24^. In this technique, steered MD (SMD) simulations are used in a wide-sampling approach to perform dozens to thousands of “pulling experiments”, in an analogy to what is done experimentally using atomic force microscopes (AFM). Allied to AFM-based SMFS, SMD has been successfully used to investigate a myriad of mechanically relevant biomolecular systems, including avidin:biotin ^25–27^, titin ^28^, human fibronectin ^28^, aquaporins ^29^, among others.

The development of an *in silico* SMFS methodology, allowed us to go even further and to fine-tune mechanical properties of protein folds ^30^. Besides protein design, our methodology allowed us to discover ultrastable protein complexes, and to decipher their intricate mechanostability mechanisms ^22,31,32^. Among these ultrastable protein complexes, the ones formed by *Staphylococci* bacteria when adhering to humans are particularly interesting ^33^. These bacteria adhere to their hosts through proteins called adhesins (Dufrêne and Viljoen 2020). Adhesins play critical roles during infection, especially during the early step of adhesion when cells are exposed to mechanical stress. During the first steps of staph infection, the interactions between adhesins and proteins of the human extracellular matrix are a key virulence factor for these bacteria ^36^, and a crucial first step of biofilm formation ^37^. These staph biofilms are associated with more than half of all nosocomial infections ^38^, with *Staphylococcus epidermidis* and *S. aureus* listed as the most common pathogens ^36,39^.

To demonstrate the advantages and limitations of AI-based protein structure prediction methods, in this perspective article we used AlphaFold to predict the structures of several *S. aureus* adhesins from the adhesion domain superfamily. First, a bioinformatics analysis was performed to select a diverse set of adhesin sequences of different *S. aureus* strains that were then used as input for AlphaFold, when structural models were generated. Then, we employed our *in silico* SMFS methodology to characterize the mechanical properties of these proteins, comparing the results to those obtained with traditional structure biology methods.

## 2 Application: Adhesin folding domains

### 2.1 How good is AlphaFold to model full length adhesins?

After selecting 48 *S. aureus* adhesins from the adhesion superfamily and we used AlphaFold 2 through the VMD’s ^40^ QwikFold plugin ^41^ batch mode to construct the models for 48 full length apo adhesin protein models. Overall, AlphaFold 2 consistently predicted the canonical folds for N2 and N3 domains for all proteins and the homologous B repeats according to each protein domain organization ^42–44^ (Fig. S1 and Table S1). Interestingly, for the collagen binding adhesin (Can), AlphaFold 2 predicted 4 additional B domains instead of the 3 described on the protein fold organization. As expected, domains such N1, the serine aspartate or fibronectin binding repeats, as well as signal sequences, were predicted as disordered.

An example of an AlphaFold prediction for the serine-aspartate repeat-containing protein E (SdrE) is shown at Figure 1. The software predicted the Ig-like N2 and N3 domains in addition to B1, B2 and B3 repeats (Fig. 1A). The N and C-ter regions normally comprise disordered regions, such as peptide signals and the SD repeats, in the case of the serine aspartate repeat proteins (Fig. 1A). A comparison between the crystal structure for SdrE (PDB ID:5WTA) containing the N2 and N3 domains and the model revealed a root mean square deviation (RMSD) of 1.31 Å for the same region (Fig. 1B), indicating that the model is a good approximation for the crystallographic structure of the Ig-like domains. This was expected since the SdrE crystal structure, among other crystals for adhesins and similar folds, were present among the structures present on AlphaFold’s training set.

**Figure 1.**
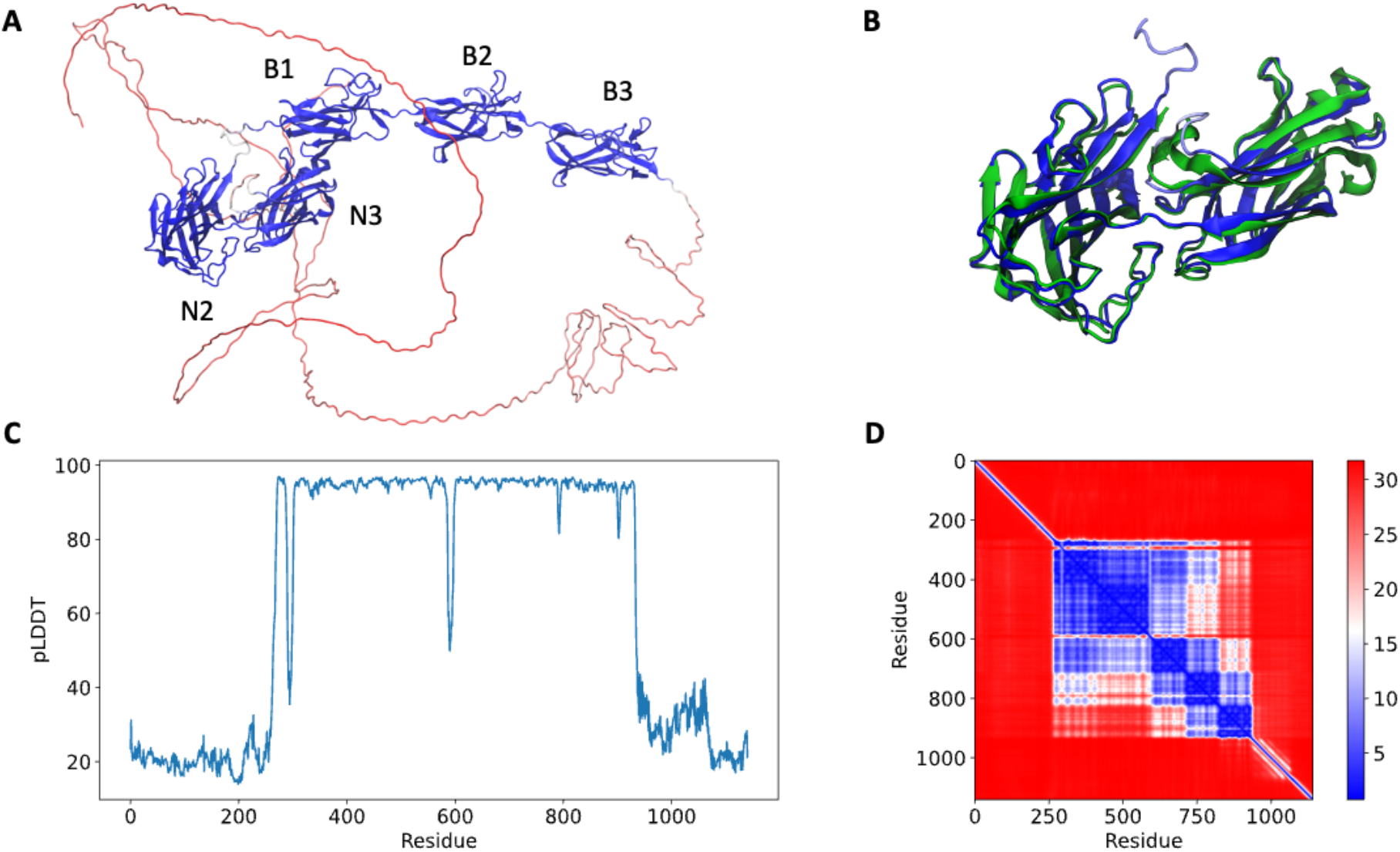
Full-length structure prediction of *S. aureus* serine-aspartate repeat protein (SdrE, Uniprot ID: Q932F7). (A) Top ranked SdrE model is represented in cartoon and it is different domains are indicated. The protein is colored by the pLDDT scores generated by AlphaFold 2 where dark blue represent regions with very high quality (pLDDT > 90) and red represent regions with very low quality (pLDDT <50). (B) Structural alignment between the N2 and N3 regions of the AlphaFold 2 model (dark blue) and SdrE crystallographic structure (cyan, PDB ID: 5WTA). (C) By residue pLDDT scores for the generated SdrE models. (D) Predicted alignment error (PAE) for the best ranked model. The color at (x, y) corresponds to the expected distance error in residue x’s position, when the prediction and true structure are aligned on residue y.

The per-residue model quality can be evaluated by pLDDT scores. In our studies, the pLDDT scores varied from ∼20 to 90 (Fig 1C) ranging from the disordered to folded regions of the proteins, which were predicted with high-quality. The confidence of the prediction can be accessed through the prediction alignment error (PAE) plots, which indicates the expected distance error in Angstrom (Fig. 1D). PAE shows low error values for the N2, N3 (big blue square) and the B domains (three small squares), corroborating the pLDDT scores for the same region and indicating high-confidence for the prediction of the mentioned domains.

### 2.2 Is AlphaFold Multimer reliable for *in-silico* force spectroscopy experiments?

*Staphylococcal* adhesins use a conserved “dock, lock, and latch” (DLL) mechanism—in which the host target, usually a peptide on the order of 15 residues, is first bound (dock), then buried (lock) between two immunoglobulin-like (Ig) fold domains N2 and N3 ^45^, and finally a “latch” connects N3 back to N2 holding the complex in place (Fig 2A). Small conformational changes on the Ig-like N2 and N3 domains could potentially impact force resilience when complexed to peptides if the DLL configuration is lost. Similar to the DLL mechanism, multiple biological phenomena rely on specific protein:protein interactions. Leveraging the initial protein structure prediction model, AlphaFold Multimer ^46^ was developed to predict structures of protein complexes for computational studies.

**Figure 2.**
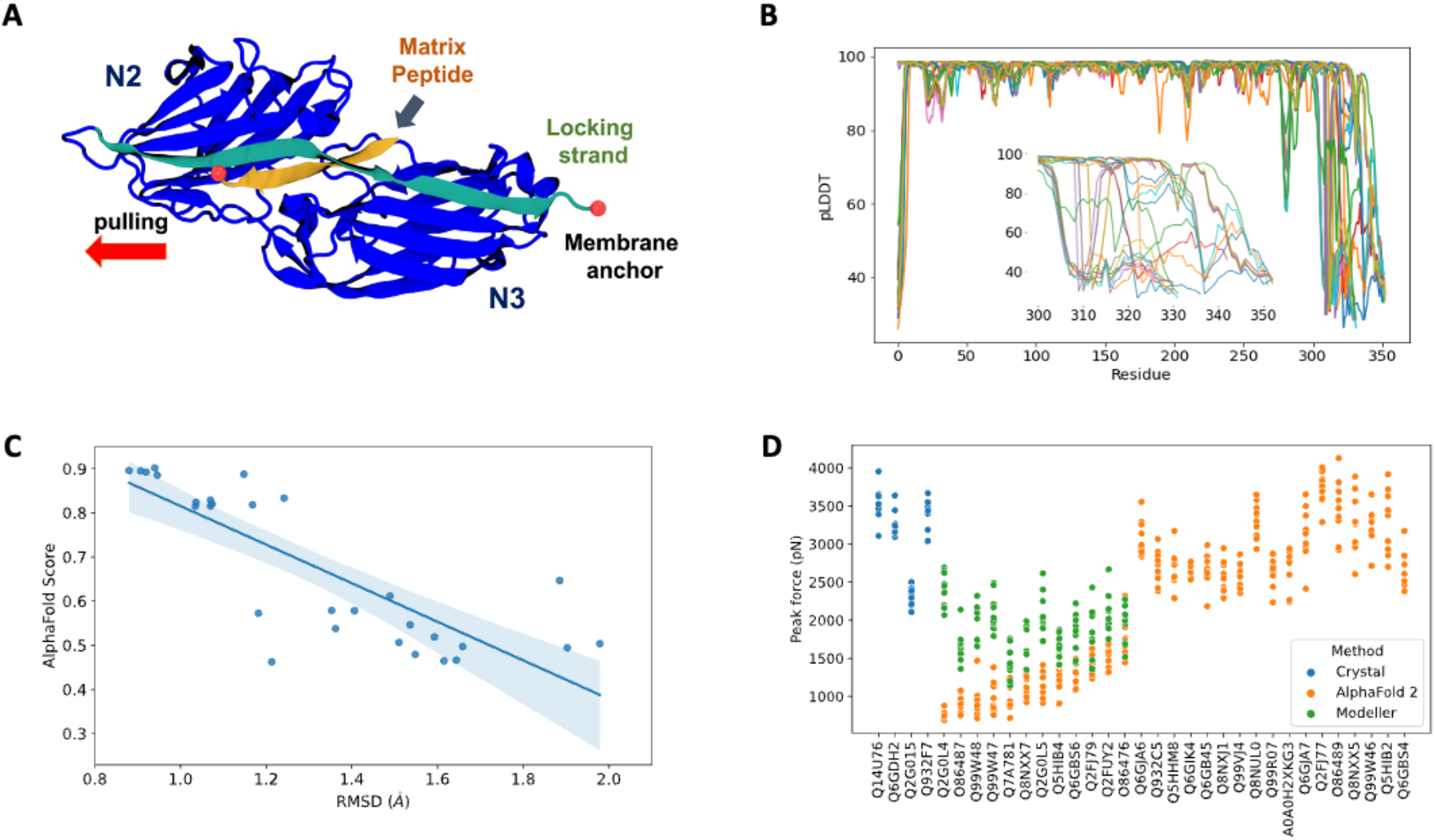
AlphaFold Multimer predictions for *S. aureus* adhesins. (A) Schematic view of adhesin’s Ig-like domain. Peptides from the host extracellular matrix are “locked” on a cleft between the N-ter N2 and N3 domains, snugly accommodated by the “locking strand”, connecting N3 to N2 by β-strand complementation (latch). SMD simulations were performed by keeping the C-ter fixed as it would be anchored to the membrane while the peptide is pulled at the opposite direction by its N-terminal. (B) By residue pLDDT scores for the top ranked model at each complex prediction. The insert shows the variation among the C-ter residues. (C) Comparison between AlphaFold Multimer score (ipTM) and RMSD values for equilibration pre-SMD simulations. (D) Peak Forces registered during SMD simulations for each studied complex. Color code indicates the origin of the departure structure: AlphaFold (orange), Modeller (green) or crystallographic (blue). Description of each accession entry are available at Table S2.

Here, we tested the reliability of *in silico* SMFS experiments performed with protein structures predicted by AlphaFold Multimer. To this end, we selected 29 adhesin sequences to be modelled in complex with extracellular matrix peptides (Table S2). AlphaFold Multimer was used to construct models for the complexes through the QwikFold ^41^ interface. Models were ranked by the predicted interface TM (ipTM) scores, used by AlphaFold Multimer, and the best ranked model for each complex was selected for SMD simulations, carried out using NAMD 3 ^47^, where the adhesins were C-terminal anchored while the peptides were pulled at a constant speed for which we measure the forces upon the dissociation of the complex. Details and parameters are described at the Supplementary Information session. As control experiments, we also initiated SMD simulations using *S. aureus* crystallographic structures of three adhesin:peptide complexes: bone sialoprotein binding protein (BBP), clumping factor A (ClfA) and SdrE.

The predicted complexes were evaluated using pLDDT scores (Fig. 2B). Most of the protein display high quality (pLDDT > 80), with exception of a very small portion of the N-terminal (10 to 15 residues) and a significant region of the C-terminal (last 50 residues, figure 2B insert). The locking strand involved on the DLL mechanism is located on the C-terminal region of the protein structure, so this loss in model quality could impact the usability of the predicted structures in high-resolution experiments such as MD or SMD simulations.

By comparing the RMSD calculated on an equilibration MD versus the general AlphaFold Multimer scores for the best ranked structures are shown at Figure 2C. We noticed that there is a correlation (Pearson correlation of 0.82, p<0.005) between the model stability and the AlphaFold Multimer scores. Therefore we can anticipate that high scored structures present less deviation from its initial configuration, suggesting a more stable or resilint fold. AlphaFold Multimer scoring is based on an interface predicted template modelling (ipTM) score that takes into account protein-protein interactions. It was shown to be more advantageous over the pTM and pLDDT scores used in AlphaFold 2 ^48^. The best ranked models on this case are a good indicator of model confidence based on the RMSD values.

After performing *in silico* SFMS experiments on all 48 complexes, we observed that the peak force profiles ranged from ∼600 to 4000 pN, a much broader range than previously simulated SdrE, BBP and ClfA complexes initiated from crystal structures (Fig. 2D). *S. aureus* adhesins are extremely mechanoresistant with rupture forces consistently on the 2000 pN regime ^49^, values which we also reproduced on this study maintaining the same *in silico* SFMS protocol used for all complexes. Considering the drastic difference in rupture forces, we found that the very low values (600-1000 pN) seen for some of the complexes might have come from inaccurate initial structures. Visual inspection of the models with low rupture forces revealed that in most cases the locking strand was modelled in an unfavorable conformation to hold the peptide in the DLL configuration, which explains the observed behavior (data not shown).

To test this hypothesis, we re-modelled those complexes using comparative modelling with Modeller ^50^ (Table S2). The models were inspected for the presence of the locking strand and simulated according to the same protocol described above (peak force profiles are shown in Figure 2D). For all cases we recover the force resilience, with peaks reaching 2000 to 3000 pN range, confirming that a high-resolution initial structure is necessary to be used for MD and SMD simulations.

## 3 Discussion

Protein structure prediction has been one of the grand challenges in Biology since the 1950’s ^51,52^. Several methods have been developed over the past 40 years that span from comparative modeling with the increase of experimentally determined protein structures by X-ray crystallography, nuclear magnetic resonance spectroscopy (NMR) and cryo-electron microscopy (cryo-EM) ^53^, but little progress was seen on *ab-initio* methodologies that rely solely on the protein sequence. But all of that changed upon the release of AlphaFold in 2021. Although AlphaFold requires only the protein sequence as input, it should not be considered an *ab-initio* method since it is built on 50 years of knowledge of protein structure determination by experimental methods. AlphaFold tremendous success took advantage of both the recent explosion of AI methods, as well as the huge database of protein structure offered by the protein data bank (PDB) ^54^.

However, as nearly any other AI-based tool, AlphaFold is biased towards its training set, meaning that the search for unusual folds is unlikely to provide an accurate result. Despite the software’s success on the folded part of most proteins, AlphaFold lacks accuracy for regions where fewer sequences are available for alignment and intrinsically disordered regions, the latter are about one third of the human proteome, present in all proteomes of all kingdoms of life, and of all viral proteomes analyzed so far ^55,56^. It also struggles with protein interfaces in homo or hetero-multimers ^46^ and other aspects of protein structures such as co-factors, post-translational modifications and DNA or RNA complexes.

In order to show how revolutionary AlphaFold is for the single-molecule biophysics community, here we put AlphaFold to the test by using it to model full length staph adhesins and estimate how stable are the protein structures. Ignoring the disordered regions, AlphaFold was able to model the Ig-like domains of adhesins as well as other key structural features of these proteins, such as the homologous B domains, for all the tested sequences. With a little refinement from *in-equilibrium* MD simulations, the generated structures could help to investigate the properties of many of the domains that still have an unknown function.

We also tested the newly developed AlphaFold Multimer to model adhesin:peptide complexes from different strains of *S. aureus* involved in biofilm formation. By comparing the force profile obtained from crystallographic structures of the complexes, we showed that AlphaFold Multimer failed to predict important key structural motifs for some of the protein complexes. Particularly, the locking strand of the adhesins, which are essential for interacting and locking the human target peptide in a tight complex with the N2 and N3 domains. It is still unclear why the predicted models worked for some cases and not for others. Limiting the set of templates to the ones where we know that the correct structures are present did not help to improve the results (data not shown). This highlights that its Multimer mode is not yet suitable to be blindly used as a peptide docking approach and the generated models should pass through a manual inspection to be suited for MD simulations.

In summary, AlphaFold is a truly revolutionary tool that is bringing a new level of structural biology to bioinformatics. Although there are many areas where its methodology can be improved, the current algorithm can be clearly employed to work alongside single-molecule biophysics experiments. It is important to note that, as any other scientific tool, particularly new ones, AlphaFold results cannot be employed blindly. Assessing the quality of the results and the usability of the predicted structures to infer function or mechanism to proteins is still the work of a trained scientist that can bring together data from multiple sources in a careful analysis of protein structure and dynamics.

## 4 Methods

### 4.1 Protein structure prediction and processing

We selected 48 *S. aureus* adhesins from the adhesion superfamily (InterPro: IPR008966) to have their full-length sequence (∼1k residues) modelled. Sequences were retrieved from the Uniprot database according to the accession numbers described on Table S1. We used AlphaFold 2 ^1^ through the VMD QwikFold plugin batch mode ^41^ to construct the models for the full length apo proteins. Current implementation of AlphaFold 2 can generate up to 24 predictions for each sequence. All models were ranked by the predicted Local Distance Difference Test (pLDDT) quality scores.

From the same protein family, we selected 29 *S. aureus* adhesins to be modelled in complex with peptides from the human extracellular matrix (Table S2). The full-length sequences retrieved from Uniprot. Before structure prediction, the sequences were trimmed to contain only the N2 and N3 domains, necessary to dock and lock the peptides. To do that, all sequences were aligned using MAFFT ^57^. HMMER ^58^ was used to generate a hidden-markov model profile using 3 *S. aureus* adhesins crystal structures that contained the Ig-like domains N2 and N3: bone sialoprotein binding protein, clumping factor B and serine-aspartate repeat-containing protein E (PDB IDs: 5CFA, 4F1Z, 5WTA, respectively). This profile was used to search against the aligned sequences and select the corresponding regions. Sequences for fibrinopeptides alfa, beta, gamma, in addition to complement factor H peptide were retrieved from available crystal structures for bone-sialoprotein binding protein (BBP), serine-aspartate repeat-containing protein G (SdrG from *S. epidermidis*), clumping factor A (ClfA) and serine-aspartate repeat-containing protein E (SdrE) (PDB IDs: 5CFA, 1R17, 2VR3, 5WTB, respectively). Proteins were later paired with each peptide based on information available on Uniprot. We used AlphaFold Multimer ^46^ through the QwikFold batch mode to construct the models. Five predictions were generated for each protein complex and ranked by the predicted interface template modelling (ipTM) scores, used by AlphaFold Multimer. The best ranked model for each complex was selected for steered molecular dynamics simulations.

Some adhesin:peptide complexes displayed very low force profiles upon SMD simulations. These were remodeled using Modeller ^50^ using as templates BBP or SdrE crystal structures. Models were generated using standard parameters and the structures followed the same protocol described at the next session.

### 4.2 Steered molecular dynamics (SMD) simulations

SMD simulations were conducted using NAMD 3 ^59^. All systems were prepared using the VMD QwikMD interface ^60,61^ where the proteins were solvated with TIP3 water model ^62^ and the total charge neutralized using NaCl 0.15 mol/L ion concentration. The CHARMM36 ^63^ force field was used to describe the system and the simulations were performed under periodic boundary conditions in the NpT ensemble with temperature maintained at 300 K using Langevin dynamics for temperature and pressure coupling, the latter kept at 1 bar. A distance cut-off of 11.0 Å was applied to short-range non-bonded interactions, whereas long-range electrostatic interactions were treated using the particle-mesh Ewald (PME) ^64^ method. Before the SMD simulations all the systems were submitted to an energy minimization protocol for 1,000 steps. Additionally, an MD simulation with position restraints in the protein backbone atoms was performed for 1 ns, with temperature ramping from 0k to 300 K in the first 0.5 ns, which served to pre-equilibrate the system. For SMD, adhesins were C-terminal anchored, while peptides were pulled at a constant speed of 5×10-5 Å/ps with a 5 kcal/mol/Å^2^ spring constant for 10 ns. Production runs were generated in ten replicas for each complex. We also selected 3 *S. aureus* crystallographic structures of adhesin:peptide complexes: BBP, ClfA and SdrE (PDB IDs: 5CFA, 2VR3 and 5WTB) to be simulated as control. Root mean square deviation (RMSD) values were calculated for the equilibrium simulation pre-SMD, for all systems. Protein images were generated using VMD; plots were rendered using python scripts.

## 6 Author Contributions

PSFG contributed to performing all the simulations, analysing data and writing of the manuscript. DEBG contributed to analysing data and discussion on AI-based methods. RCB coordinated the project, contributed to writing and discussion on *in silico* force spectroscopy, proof-reading, manuscript revision and approval of the submitted version.

## 8 Acknowledgments

We thank Auburn University and the College of Sciences and Mathematics for the computational resources provided by Dr. Bernardi faculty startup funds. We thank Dr. Marcelo Melo for the fruitful discussions.

## 9 Data Availability Statement

The data that support the findings of this study are available from the corresponding author, Dr. Bernardi, upon reasonable request.

## References

1. Jumper, J. et al. Highly accurate protein structure prediction with AlphaFold. Nature 2021 596:7873 596, 583–589 (2021).

2. Callaway, E. What’s next for AlphaFold and the AI protein-folding revolution. Nature 604, 234–238 (2022).

3. Baek, M. et al. Accurate prediction of protein structures and interactions using a three-track neural network. Science 373, 871–876 (2021).

4. Tunyasuvunakool, K. et al. Highly accurate protein structure prediction for the human proteome. Nature 596, 590–596 (2021).

5. Jumper, J. et al. Applying and improving AlphaFold at CASP14. Proteins: Structure, Function, and Bioinformatics 89, 1711–1721 (2021).

6. Varadi, M. et al. AlphaFold Protein Structure Database: massively expanding the structural coverage of protein-sequence space with high-accuracy models. Nucleic Acids Research 50, D439–D444 (2022).

7. Hartmann, S. et al. Structure and Protein-Protein Interactions of Ice Nucleation Proteins Drive Their Activity. bioRxiv 2022.01.21.477219 (2022) doi:10.1101/2022.01.21.477219.

8. Mosalaganti, S. et al. Artificial intelligence reveals nuclear pore complexity. BioRxiv 2021.10.26.465776 (2021) doi:10.1101/2021.10.26.465776.

9. Skolnick, J., Gao, M., Zhou, H. & Singh, S. AlphaFold 2: Why It Works and Its Implications for Understanding the Relationships of Protein Sequence, Structure, and Function. Journal of Chemical Information and Modeling 61, 4827–4831 (2021).

10. Perrakis, A. & Sixma, T. K. AI revolutions in biology. EMBO Rep 22, e54046 (2021).

11. Outeiral, C., Nissley, D. A. & Deane, C. M. Current structure predictors are not learning the physics of protein folding. Bioinformatics 38, 1881–1887 (2022).

12. Eisenstein, M. Artificial intelligence powers protein-folding predictions. Nature 599, 706–708 (2021).

13. Akdel, M. et al. A structural biology community assessment of AlphaFold 2 applications. bioRxiv 2021.09.26.461876 (2021) doi:10.1101/2021.09.26.461876.

14. Mullard, A. What does AlphaFold mean for drug discovery? Nat Rev Drug Discov 20, 725–727 (2021).

15. Buel, G. R. & Walters, K. J. Can AlphaFold2 predict the impact of missense mutations on structure? Nat Struct Mol Biol 29, (2022).

16. Lee, E. H., Hsin, J., Sotomayor, M., Comellas, G. & Schulten, K. Discovery through the computational microscope. Structure 17, 1295–1306 (2009).

17. Perilla, J. R. et al. Molecular dynamics simulations of large macromolecular complexes. Curr Opin Struct Biol 31, 64–74 (2015).

18. Dror, R. O., Dirks, R. M., Grossman, J. P., Xu, H. & Shaw, D. E. Biomolecular simulation: a computational microscope for molecular biology. Annu Rev Biophys 41, 429–452 (2012).

19. Heo, L. & Feig, M. Experimental accuracy in protein structure refinement via molecular dynamics simulations. Proc Natl Acad Sci U S A 115, 13276–13281 (2018).

20. Melo, M. C. R. et al. NAMD goes quantum: An integrative suite for hybrid simulations. Nature Methods 15, 351–354 (2018).

21. Bernardi, R. C. & Pascutti, P. G. Hybrid QM/MM Molecular Dynamics Study of Benzocaine in a Membrane Environment: How Does a Quantum Mechanical Treatment of Both Anesthetic and Lipids Affect Their Interaction. Journal of Chemical Theory and Computation 8, 2197–2203 (2012).

22. Bernardi, R. C. et al. Mechanisms of Nanonewton Mechanostability in a Protein Complex Revealed by Molecular Dynamics Simulations and Single-Molecule Force Spectroscopy. J Am Chem Soc 141, 14752–14763 (2019).

23. Sedlak, S. M. et al. Direction Matters: Monovalent Streptavidin/Biotin Complex under Load. Nano Letters 19, 3415–3421 (2019).

24. Sedlak, S. M., Schendel, L. C., Gaub, H. E. & Bernardi, R. C. Streptavidin/biotin: Tethering geometry defines unbinding mechanics. Science Advances 6, (2020).

25. Izrailev, S., Stepaniants, S., Balsera, M., Oono, Y. & Schulten, K. Molecular dynamics study of unbinding of the avidin-biotin complex. Biophysical Journal 72, 1568 (1997).

26. Grubmüller, H., Heymann, B. & Tavan, P. Ligand Binding: Molecular Mechanics Calculation of the Streptavidin-Biotin Rupture Force. Science (1979) 271, 997–999 (1996).

27. Merkel, R., Nassoy, P., Leung, A., Ritchie, K. & Evans, E. Energy landscapes of receptor-ligand bonds explored with dynamic force spectroscopy. Nature 397, 50–53 (1999).

28. Gao, M., Craig, D., Vogel, V. & Schulten, K. Identifying Unfolding Intermediates of FN-III10 by Steered Molecular Dynamics. Journal of Molecular Biology 323, 939–950 (2002).

29. de Groot, B. L., Hub, J. S. & Grubmüller, H. Dynamics and energetics of permeation through aquaporins. What Do we learn from molecular dynamics simulations? Handbook of Experimental Pharmacology 190, 57–76 (2009).

30. Verdorfer, T. et al. Combining in Vitro and in Silico Single-Molecule Force Spectroscopy to Characterize and Tune Cellulosomal Scaffoldin Mechanics. J Am Chem Soc 139, 17841–17852 (2017).

31. Schoeler, C. et al. Ultrastable cellulosome-adhesion complex tightens under load. Nature Communications 2014 5:1 5, 1–8 (2014).

32. Liu, Z. et al. High force catch bond mechanism of bacterial adhesion in the human gut. Nature Communications 2020 11:1 11, 1–12 (2020).

33. Herman-Bausier, P. & Dufrêne, Y. F. Force matters in hospital-acquired infections. Science (1979) 359, 1464–1465 (2018).

34. Dufrêne, Y. F. & Viljoen, A. Binding Strength of Gram-Positive Bacterial Adhesins. Frontiers in Microbiology 11, 1457 (2020).

35. TJ, F. & M, H. Surface protein adhesins of Staphylococcus aureus. Trends Microbiol 6, 484–488 (1998).

36. Otto, M. Staphylococcal biofilms. Current Topics in Microbiology and Immunology vol. 322 207–228 (2008).

37. Latasa, C., Solano, C., Penadés, J. R. & Lasa, I. Biofilm-associated proteins. Comptes Rendus - Biologies vol. 329 849–857 (2006).

38. Jamal, M. et al. Bacterial biofilm and associated infections. J Chin Med Assoc 81, 7–11 (2018).

39. Schilcher, K. & Horswill, A. R. Staphylococcal Biofilm Development: Structure, Regulation, and Treatment Strategies. Microbiology and Molecular Biology Reviews 84, (2020).

40. Humphrey, W., Dalke, A. & Schulten, K. VMD: Visual molecular dynamics. Journal of Molecular Graphics 14, 33–38 (1996).

41. Gomes, D. E. B., da Silva Figueiredo Celestino Gomes, P. & C Bernardi, R. QwikMD 2.0: bridging the gap between sequence, structure, and protein function. Biophysical Journal 121, 132a (2022).

42. Foster, T. J. & Höök, M. Surface protein adhesins of Staphylococcus aureus. Trends in Microbiology 6, 484–488 (1998).

43. Ganesh, V. K. et al. Structural and biochemical characterization of Staphylococcus aureus clumping factor B/ligand interactions. Journal of Biological Chemistry 286, 25963–25972 (2011).

44. Foster, T. J., Geoghegan, J. A., Ganesh, V. K. & Höök, M. Adhesion, invasion and evasion: the many functions of the surface proteins of Staphylococcus aureus. Nature Reviews Microbiology 2013 12:1 12, 49–62 (2013).

45. Ponnuraj, K. et al. A “dock, lock, and latch”Structural Model for a Staphylococcal Adhesin Binding to Fibrinogen. Cell 115, 217–228 (2003).

46. Evans, R. et al. Protein complex prediction with AlphaFold-Multimer. BioRxiv 2021.10.04.463034 (2022) doi:10.1101/2021.10.04.463034.

47. Phillips, J. C. et al. Scalable molecular dynamics on CPU and GPU architectures with NAMD. The Journal of Chemical Physics 153, 44130 (2020).

48. Gao, M., Nakajima An, D., Parks, J. M. & Skolnick, J. AF2Complex predicts direct physical interactions in multimeric proteins with deep learning. Nature Communications 2022 13:1 13, 1–13 (2022).

49. Milles, L. F., Schulten, K., Gaub, H. E. & Bernardi, R. C. Molecular mechanism of extreme mechanostability in a pathogen adhesin. Science (1979) 359, 1527–1533 (2018).

50. Eswar, N., Eramian, D., Webb, B., Shen, M.-Y. & Sali, A. Protein structure modeling with MODELLER. Methods Mol Biol 426, 145–59 (2008).

51. Dill, K. A. & MacCallum, J. L. The protein-folding problem, 50 years on. Science 338, 1042–1046 (2012).

52. Dill, K. A., Ozkan, S. B., Shell, M. S. & Weikl, T. R. The Protein Folding Problem. Annu Rev Biophys 37, 289 (2008).

53. Goh, B. C. et al. Computational Methodologies for Real-Space Structural Refinement of Large Macromolecular Complexes. http://dx.doi.org/10.1146/annurev-biophys-062215-011113 45, 253–278 (2016).

54. Berman, H. M. et al. The Protein Data Bank. Nucleic Acids Res 28, 235–242 (2000).

55. Peng, Z. et al. Exceptionally abundant exceptions: comprehensive characterization of intrinsic disorder in all domains of life. Cell Mol Life Sci 72, 137–151 (2015).

56. Xue, B., Dunker, A. K. & Uversky, V. N. Orderly order in protein intrinsic disorder distribution: disorder in 3500 proteomes from viruses and the three domains of life. J Biomol Struct Dyn 30, 137–149 (2012).

57. Nakamura, T., Yamada, K. D., Tomii, K. & Katoh, K. Parallelization of MAFFT for large-scale multiple sequence alignments. Bioinformatics 34, 2490–2492 (2018).

58. Eddy, S. R. Accelerated Profile HMM Searches. PLOS Computational Biology 7, e1002195 (2011).

59. Phillips, J. C. et al. Scalable molecular dynamics on CPU and GPU architectures with NAMD. The Journal of Chemical Physics 153, 044130 (2020).

60. Humphrey, W., Dalke, A. & Schulten, K. VMD: Visual molecular dynamics. Journal of Molecular Graphics 14, 33–38 (1996).

61. Ribeiro, J. v. et al. QwikMD — Integrative Molecular Dynamics Toolkit for Novices and Experts. Scientific Reports 2016 6:1 6, 1–14 (2016).

62. Jorgensen, W. L. & Jenson, C. Temperature dependence of TIP3P, SPC, and TIP4P water from NPT Monte Carlo simulations: Seeking temperatures of maximum density. Journal of Computational Chemistry 19, 1179–1186 (1998).

63. Best, R. B. et al. Optimization of the additive CHARMM all-atom protein force field targeting improved sampling of the backbone φ, ψ and side-chain χ1 and χ2 Dihedral Angles. Journal of Chemical Theory and Computation 8, 3257–3273 (2012).

64. Darden, T., York, D. & Pedersen, L. Particle mesh Ewald: An N⋅log(N) method for Ewald sums in large systems. The Journal of Chemical Physics 98, 10089 (1993).

