## Supplementary Material for "Protein structure prediction in the era of AI: challenges and limitations when applying to *in-silico* force spectroscopy"

### 1 Supplementary figures

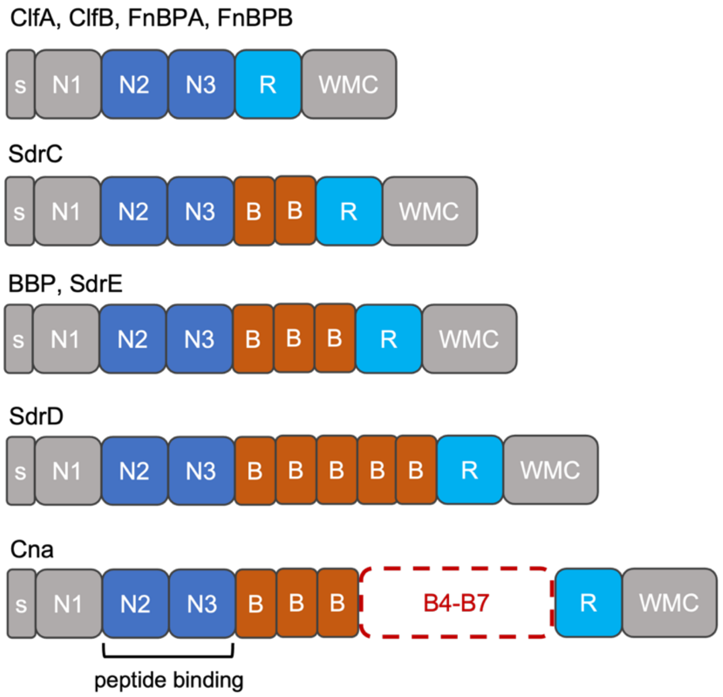


**Supplementary Figure 1.** *S. aureus* adhesin domain organization for clumping factor A (ClfA), Clumping factor B (ClfB), Fibronecting binding protein A (FnBPA), Fibronecting binding protein B (FnBPB), Serine-aspartate repeat protein C (SdrC), Serine-aspartate repeat protein D (SdrD), Serine-aspartate repeat protein E (SdrE), Bone sialoprotein binding protein (BBP) and Collagen binding adhesin (Cna). S, signal peptide; N-ter domains N1 to N3; B, homologous repeats; R, serine-aspartate or fibronectin binding repeats; W, wall-spanning region; M, membrane anchor; C, cytoplasmic tail. The ligand binding region is formed in a cleft between the N2 and N3 domains. Regions colored in gray had no defined structure predicted by AlphaFold 2. The insert displayed as dashed lines represents additional B domains predicted for Cna.

### 2 Supplementary Tables

Table S1: *S. aureus* adhesins modelled by AlphaFold 2. Full length sequences were obtained from Uniprot according to the described entries. Predictions for N2/N3 are described by a "yes" and the number of modelled homologous B domains are also described. Failed AlphaFold 2 predictions are highlighted in bold.

| Uniprot ID | Name | N2/N3 | B |
|---|---|---|---|
| Q14U76 | Bone sialoprotein-binding protein | y | 3 |
| Q6GJA6 | Bone sialoprotein-binding protein | y | 3 |
| Q2G015 | Clumping factor A | y | 0 |
| Q53653 | Clumping factor A | y | 0 |
| Q5HHM8 | Clumping factor A | y | 0 |
| Q6GB45 | Clumping factor A | y | 0 |
| Q6GIK4 | Clumping factor A | y | 0 |
| Q8NXJ1 | Clumping factor A | y | 0 |
| Q932C5 | Clumping factor A | y | 0 |
| Q99VJ4 | Clumping factor A | y | 0 |
| O86476 | Clumping factor B | y | 0 |
| Q2FUY2 | Clumping factor B | y | 0 |
| Q5HCR7 | Clumping factor B | y | 0 |
| Q6G644 | Clumping factor B | y | 0 |
| Q6GDH2 | Clumping factor B | y | 0 |
| Q7A382 | Clumping factor B | y | 0 |
| Q8NUL0 | Clumping factor B | y | 0 |
| Q99R07 | Clumping factor B | y | 0 |
| Q53654 | Collagen adhesin | y | 7 |
| A7X6I5 | Fibronectin-binding protein A | y | 0 |
| P14738 | Fibronectin-binding protein A | y | 0 |
| Q2FE03 | Fibronectin-binding protein A | y | 0 |
| Q2YW62 | Fibronectin-binding protein A | y | 0 |
| Q5HD51 | Fibronectin-binding protein A | y | 0 |
| Q6G6H3 | Fibronectin-binding protein A | y | 0 |
| Q6GDU5 | Fibronectin-binding protein A | y | 0 |
| **Q7A3J7** | Fibronectin-binding protein A | | |
| Q8NUU7 | Fibronectin-binding protein A | y | 0 |
| Q99RD2 | Fibronectin-binding protein A | y | 0 |
| A0A0H2XKG3 | Fibronectin-binding protein B | y | 0 |
| O86487 | Serine-aspartate repeat-containing protein C | y | 2 |
| Q2FJ79 | Serine-aspartate repeat-containing protein C | y | 2 |
| Q2G0L5 | Serine-aspartate repeat-containing protein C | y | 2 |
| Q5HIB4 | Serine-aspartate repeat-containing protein C | y | 2 |

| | | | |
|---|---|---|---|
| Q6GBS6 | Serine-aspartate repeat-containing protein C | y | 2 |
| Q6GJA7 | Serine-aspartate repeat-containing protein C | y | 2 |
| Q7A781 | Serine-aspartate repeat-containing protein C | y | 2 |
| Q8NXX7 | Serine-aspartate repeat-containing protein C | y | 2 |
| Q99W48 | Serine-aspartate repeat-containing protein C | y | 2 |
| **O86488** | Serine-aspartate repeat-containing protein D | | |
| Q2FJ78 | Serine-aspartate repeat-containing protein D | y | 5 |
| **Q2G0L4** | Serine-aspartate repeat-containing protein D | | |
| **Q5HIB3** | Serine-aspartate repeat-containing protein D | | |
| **Q6GBS5** | Serine-aspartate repeat-containing protein D | | |
| Q7A780 | Serine-aspartate repeat-containing protein D | y | 5 |
| **Q8NXX6** | Serine-aspartate repeat-containing protein D | | |
| Q99W47 | Serine-aspartate repeat-containing protein D | y | 5 |
| O86489 | Serine-aspartate repeat-containing protein E | y | 3 |
| Q2FJ77 | Serine-aspartate repeat-containing protein E | y | 3 |
| Q5HIB2 | Serine-aspartate repeat-containing protein E | y | 3 |
| Q6GBS4 | Serine-aspartate repeat-containing protein E | y | 3 |
| Q8NXX5 | Serine-aspartate repeat-containing protein E | y | 3 |
| Q932F7 | Serine-aspartate repeat-containing protein E | y | 3 |
| Q99W46 | Serine-aspartate repeat-containing protein E | y | 3 |

Table S2: *S. aureus* adhesins and the respective peptides complexes used on the AlphaFold Multimer predictions. Entries are organized by solving method; the ones marked as solved by Modeller were updated after obtention of low force profiles when using the predicted AlphaFold Multimer structures. The full length sequences were obtained according to their Uniprot ID as described below and post-processed to contain only the N2-N3 domains as described at the Methods section below.

| Solving method | Uniprot ID | Protein | ligand | Strain |
|---|---|---|---|---|
| X-ray (PDB ID:5CFA) | Q14U76 | Bone sialoprotein-binding protein | Fibrinogen alpha | not informed |
| X-ray (PDB ID:2VR3) | Q2G015 | Clumping factor A | Fibrinogen gamma | strain NCTC 8325 / PS 47 |
| X-ray (PDB ID: 5WTB) | Q932F7 | Serine-aspartate repeat-containing protein E | Complement factor H | Mu50 / ATCC 700699 |
| AlphaFold | Q6GJA6 | Bone sialoprotein-binding protein | Fibrinogen alpha | MRSA252 |
| AlphaFold | Q932C5 | Clumping factor A | Fibrinogen gamma | Mu50 / ATCC 700699 |
| AlphaFold | Q5HHM8 | Clumping factor A | Fibrinogen gamma | COL |
| AlphaFold | Q6GIK4 | Clumping factor A | Fibrinogen gamma | MRSA252 |
| AlphaFold | Q6GB45 | Clumping factor A | Fibrinogen gamma | MSSA476 |
| AlphaFold | Q8NXJ1 | Clumping factor A | Fibrinogen gamma | MW2 |
| AlphaFold | Q99VJ4 | Clumping factor A | Fibrinogen gamma | N315 |
| AlphaFold | Q8NUL0 | Clumping factor B | Fibrinogen alpha | MW2 |
| AlphaFold | Q99R07 | Clumping factor B | Fibrinogen beta | Mu50 / ATCC 700699 |
| AlphaFold | A0A0H2XKG3 | Fibronectin-binding protein B | Fibrinogen beta | USA300 |
| AlphaFold | Q6GJA7 | Serine-aspartate repeat-containing protein C | Fibrinogen alpha | MRSA252 |
| AlphaFold | Q2FJ77 | Serine-aspartate repeat-containing protein E | Complement factor H | USA300 |
| AlphaFold | O86489 | Serine-aspartate repeat-containing protein E | Complement factor H | Newman |

| | | | | |
|---|---|---|---|---|
| AlphaFold | Q8NXX5 | Serine-aspartate repeat-containing protein E | Complement factor H | MW2 |
| AlphaFold | Q99W46 | Serine-aspartate repeat-containing protein E | Complement factor H | N315 |
| AlphaFold | Q5HIB2 | Serine-aspartate repeat-containing protein E | Complement factor H | COL |
| AlphaFold | Q6GBS4 | Serine-aspartate repeat-containing protein E | Complement factor H | MSSA476 |
| Modeller | Q2FUY2 | Clumping factor B | Fibrinogen alpha | NCTC 8325 / PS 47 |
| Modeller | O86476 | Clumping factor B | Fibrinogen alpha | Newman |
| Modeller | Q2G0L5 | Serine-aspartate repeat-containing protein C | Fibrinogen alpha | NCTC 8325 / PS 47 |
| Modeller | Q99W48 | Serine-aspartate repeat-containing protein C | Complement factor H | Mu50 / ATCC 700699 |
| Modeller | Q6GBS6 | Serine-aspartate repeat-containing protein C | Fibrinogen alpha | MSSA476 |
| Modeller | Q2FJ79 | Serine-aspartate repeat-containing protein C | Fibrinogen alpha | USA300 |
| Modeller | Q8NXX7 | Serine-aspartate repeat-containing protein C | Fibrinogen alpha | MW2 |
| Modeller | Q5HIB4 | Serine-aspartate repeat-containing protein C | Fibrinogen alpha | COL |
| Modeller | O86487 | Serine-aspartate repeat-containing protein C | Fibrinogen alpha | Newman |
| Modeller | Q7A781 | Serine-aspartate repeat-containing protein C | Fibrinogen alpha | N315 |
| Modeller | Q2G0L4 | Serine-aspartate repeat-containing protein D | Fibrinogen alpha | NCTC 8325 / PS 47 |
| Modeller | Q99W47 | Serine-aspartate repeat-containing protein D | Complement factor H | Mu50 / ATCC 700699 |
